# Critical Scaling Laws and Universality Classes in Biomolecular Condensates

**DOI:** 10.64898/2026.06.24.734243

**Authors:** Haoyu Song, Guorong Hu, Xiangyu Wu, Xialan Zhang, Jingyuan Li

**Affiliations:** School of Physics, Zhejiang University, Hangzhou 310058, PR China; Institute for Advanced Study in Physics, Zhejiang University, Hangzhou 310058, PR China

## Abstract

Biomolecular condensates are widespread cellular self-assembled structures with essential functions. There are suggestions of condensates formed by different proteins being near criticality. However, systematic investigation of the criticality of condensates is absent, and critical exponents defining their universality class have not been found. Here, using long-time simulations, we show that condensates exhibit typical critical phenomena, including scale-free spatiotemporal correlations, critical slowing down, divergence of correlation length and dynamic scaling. From these scaling behaviors, a set of critical exponents is determined. Based on dynamic critical exponent, diverse condensates can be divided into two distinct universality classes, arising from differences in their molecular components and interaction types.

---

The notion that living systems operate near the critical point of a phase transition has long been compelling in biophysics [1–3]. Numerous biological systems, such as flocking birds [4,5], the brain [3,6], cell clusters [7,8], and gene regulatory networks [9,10], are regarded as critical systems. The properties of criticality, including scale-free correlations and a trade-off between susceptibility and robustness, provide significant functional advantages to these systems. For instance, scale-free correlations enable birds to fly collectively as a whole flock, regardless of flock size [4,11]. The brain can respond sensitively to external stimuli while maintaining resilience against small perturbations [12,13], like damage of a subset of neurons. The study of biological criticality may help uncover the universal principles underlying the organization and operation of life.

Biomolecular condensates, a type of self-assembled structures in living cells, have recently been recognized as membraneless organelles that spatially and temporally organize cellular functions [14–16]. The condensates spontaneously form via liquid-liquid phase transition, driven by multivalent interactions among biomolecules [17,18]. Growing evidence suggests that condensates appear to be near criticality [19–22]. For instance, the surface tension of condensates displays a power-law dependence with respect to reduced temperature [19]. Condensates such as nucleoli follow a broad power-law size distribution [22]. Additionally, the formation and disassembly of condensates respond sensitively to stimuli, including environmental changes [23,24]. Despite these observations, there is still a lack of systematic investigations on the statistical-physical hallmarks of criticality in biomolecular condensates. Furthermore, whether diverse condensates can be characterized by a set of critical exponents and assigned to a specific universality class remains a key unresolved question.

Here, we performed a statistical analysis of critical behaviors and critical exponents in a typical condensate system (FUS-PrLD condensate), and further explored the universality across various condensates, using long-time molecular dynamics (MD) simulations. FUS-PrLD is an intrinsically disordered protein (IDP) which can form condensate *in vitro* and assemble into membraneless organelles *in vivo*, e.g., stress granules [25,26]. In this condensate, we found scale-free spatial and temporal correlations, manifested as a power-law structure factor *S*(*q*) and the 1/*f* power spectrum. Furthermore, we found that the condensate exhibits critical slowing down with respect to reduced temperature *t*, i.e., its relaxation time scaling as *τ* ∼ *t*^−*vz*^ (*vz* = 0.93), and the divergence of correlation length *ξ* ∼ *t*^−*v*^ (*v* = 0.26). We also identified the dynamic scaling law *τ* ∼ *ξ*^*z*^, with a dynamic critical exponent *z* = 3.6. Notably, these critical exponents satisfy the theoretical relationship *τ* ∼ *ξ*^*z*^ ∼ (*t*^−*v*^)^*z*^ ∼ *t*^−*vz*^. Finally, we extended the study to various condensate systems and found that they can be divided into two classes based on dynamic critical exponent *z*: condensates composed solely of IDPs exhibit *z* ≈ 3.6, whereas those containing structural domains show *z* ≈ 2.6. This distinction in universality classes arises from the different interaction types within condensates, which are influenced by the molecular features of their components (i.e., whether containing structural domains). Taken together, our results demonstrate that condensates are critical systems which can be characterized by specific critical exponents, and belong to two distinct universality classes.

Fig. 1(a) shows a snapshot of the FUS-PrLD condensate from our all-atom simulations (see the Supplemental Material for simulation details). The internal structure of the condensate shows nanoscale heterogeneity, with regions of sparse or dense residue density. It is consistent with the experimental reports of structure heterogeneity and nanoclusters in condensates [27]. The local residue density fluctuation over time (i.e., residues within a radius of 4 nm around the center of mass of the condensate) was then computed [see Fig. 1(b) inset]. The rapid density fluctuation reflects the highly dynamic nature of the condensate, in accord with its liquid-like feature. We further calculated the power spectrum *S*(*f*) of density fluctuation as follows:

**FIG. 1.**
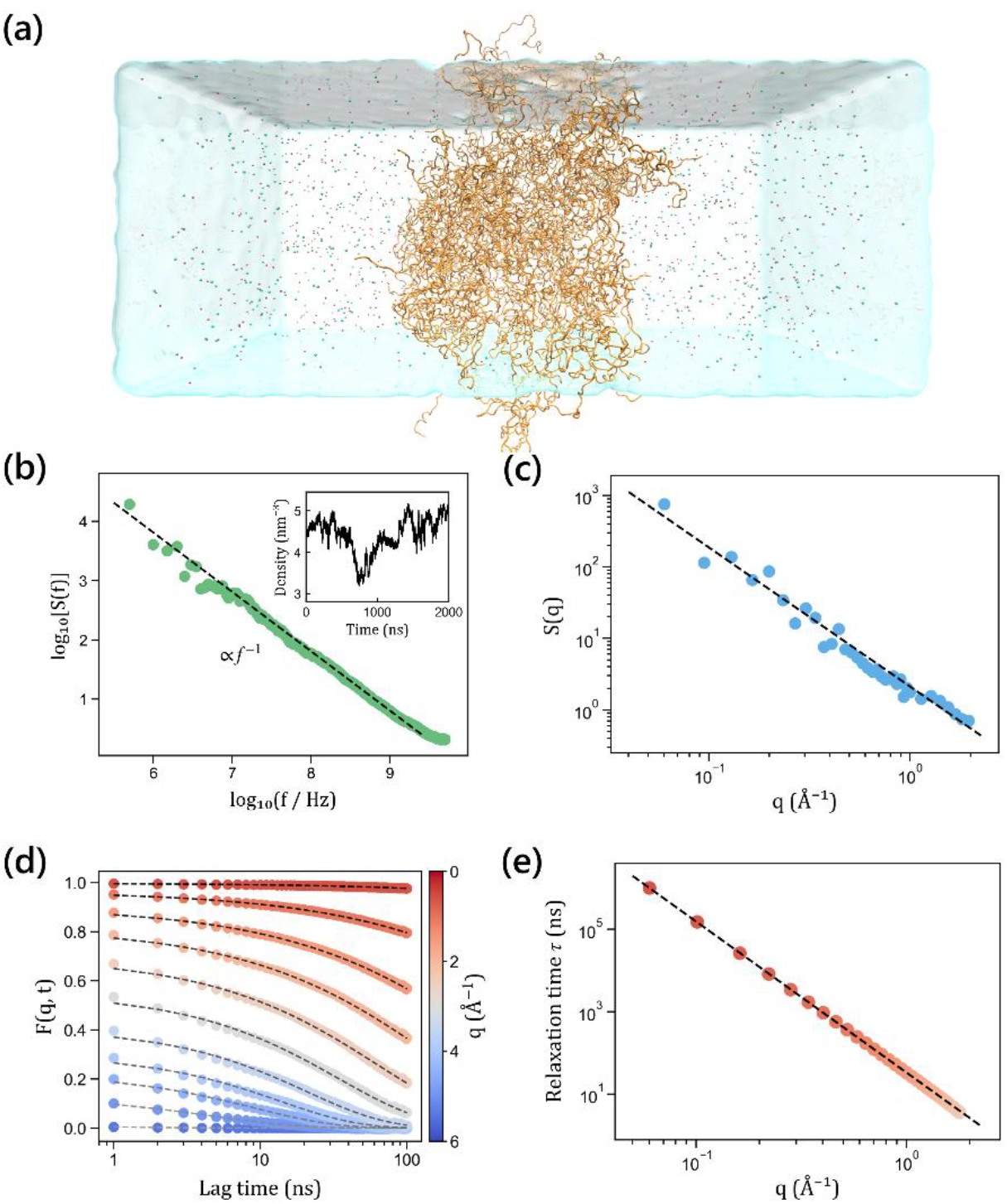
Biomolecular condensate and its power-law behavior in space and time. (a) Representative snapshot of the FUS-PrLD condensate (orange) in a slab geometry, with Na^+^ ions (red spheres), Cl^−^ ions (green spheres), and water (light blue). (b) The 1/*f* noise in the structure fluctuations of the condensate. (Inset) Time evolution of the large-scale density fluctuations of residues within a 4-nm radius of the condensate’s center of mass. (c) Power-law structure factor *S*(*q*) of the condensate. (d) Decay of the intermediate scattering factor (ISF), *F*(*q, t*), with lag time for various wavenumbers *q*. (e) Power-law dynamics between spatial scale *q* and corresponding relaxation time *τ* within the FUS-PrLD condensate.

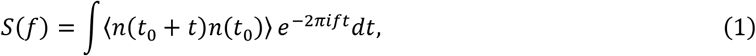

where *n* is the residue density and ⟨ · ⟩ denotes an average over all times *t*_0_. The power spectrum scales as *f*^−1^ [Fig. 1(b)], commonly referred to as “1/*f* noise” [28], indicating scale-free temporal correlation in the condensate’s conformational dynamics. To further characterize the condensate structure, we analyzed its static structure factor *S*(*q*) [see Fig. 1(c)]. *S*(*q*) of the FUS-PrLD condensate exhibits a power-law scaling with wavenumber *q*, consistent with small-angle X-ray scattering measurements on other condensates [29]. The power-law *S*(*q*) suggests scale-free spatial correlation within the condensate.

It has been argued that scaling in space and time need not be related in some systems [30]. To examine whether the observed spatial and temporal scaling behaviors in the FUS-PrLD condensate are linked, we introduced time into static *S*(*q*) to calculate the intermediate scattering factor (ISF) *F*(*q, t*) as

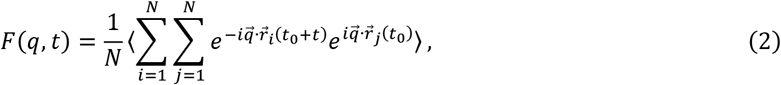

where 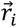 is position of the *i*-th atom and *N* is the total number of atoms in the condensate. As shown in Fig. 1(d), the decay of the *F*(*q, t*) curves become slower with increasing spatial scale (i.e., decreasing *q*). By fitting the curves with stretched exponential [red dashed lines in Fig. 1(d)], we obtained the relaxation time *τ* for each *q* [31]. Fig. 1(e) reveals a power‐law coupling between relaxation time and wavenumber:

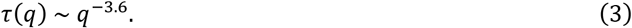

So far, we found scale-free spatiotemporal correlations and a power-law dynamic scaling between space and time in the conformation dynamics of the FUS-PrLD condensate, which are hallmarks of a critical system [31–33]. It should be noted that if condensate is indeed at criticality, the spatial range over which this dynamic scaling holds should extend as the system size increases, due to the scale-free nature of the system — a point that will be discussed later.

It is well acknowledged that the physical variables of a system near criticality obey scaling laws with respect to reduced temperature *t* = *T*/*T*_*c*_ − 1, from which the critical exponents can be derived. Previous studies have shown that the formation of condensates can be modulated by temperature, and the FUS-PrLD condensate generally forms near room temperature *in vitro* [34]. Therefore, we further investigated the temperature dependence of FUS-PrLD condensate behavior by performing coarse-grained simulations across a temperature gradient (see details in the Supplemental Material), with an estimated *T*_*c*_ ≈ 293 *K* (see Fig. S11). The overall relaxation time *τ*_*sys*_ of the condensate, defined as *τ* at the minimum wavenumber *q*_*min*_ corresponding to the system size, was determined from the ISF curves at each temperature. The relationship between *τ*_*sys*_ and reduced temperature *t* is shown in Fig. 2(a). Consistent with theoretical expectations [35], *τ*_*sys*_ follows a power-law dependence on *t*:

**FIG. 2.**
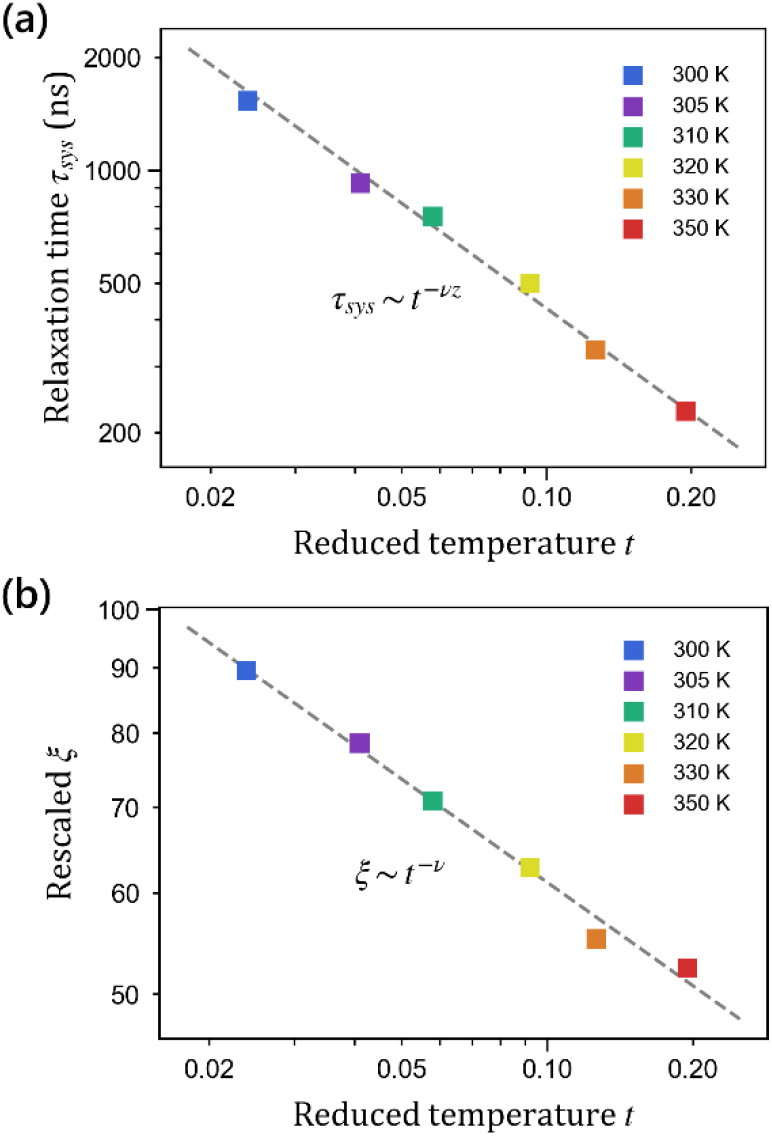
Scaling laws of the condensate with respect to reduced temperature. (a) Critical slowing down. The relaxation time of the condensate at different temperatures follows a power-law relation *τ*_*sys*_ ∼ *t*^−*vz*^, with a fitted exponent *vz* = 0.93 (dashed grey line). (b) Divergence of the correlation length. The correlation length of the condensate scales as *ξ* ∼ *t*^−*v*^, with *v* = 0.26.

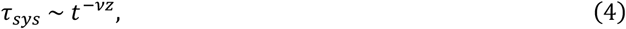

where *vz* is a combination of critical exponents, with a fitted value of *vz* = 0.93. This relationship indicates that the condensate exhibits “critical slowing down”, meaning that the relaxation dynamics of a system become slower as the critical point is approached.

According to the theory of critical phenomena, the combined exponent *vz* is a composite of two fundamental exponents: *vz* = *v* · *z*, where *v* governs the divergence of the correlation length *ξ* near criticality, and *z* is the dynamic critical exponent [35]. To further determine the critical exponents of the condensate system, we then aimed to study these two critical exponents separately. We first sought to determine *v* by estimating the correlation length *ξ* of FUS-PrLD condensates at different temperatures (see details in the Supplemental Material). As shown in Fig. 2(b), *ξ* gradually increases as temperature decreases. Moreover, *ξ* also follows a power-law dependence on the reduced temperature *t*, expressed as

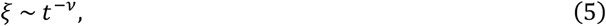

consistent with the scaling law expected in critical systems. Fitting the data yielded a value of the correlation length critical exponent *v* = 0.26.

We next investigated another critical exponent *z*. Typically, *z* can be directly obtained by plotting *τ*_*sys*_ against the corresponding *ξ* at different temperatures [see Fig. 3(a)]. The power-law fitting shows the dynamic critical exponent *z* = 3.6. This relationship reflects the dynamic scaling law *τ*_*sys*_ ∼ *ξ*^*z*^ between the condensate’s relaxation dynamics and its system size in space. It should be noted that this physical interpretation is similar to that of *τ*(*q*) discussed earlier (namely, a power‐law coupling between relaxation time and spatial scale), except that *τ*(*q*) was defined within the condensate system and was cut off by the system size. Additionally, the value of *z* is equal to the exponent in *τ*(*q*) ∼ *q*^−3.6^, where the opposite sign arises from the reciprocal relationship between real space and q-space. Thus, we proposed that the dynamic scaling law *τ*_*sys*_ ∼ *ξ*^3.6^ and the previous *τ*(*q*) ∼ *q*^−3.6^ share the same physical nature. To test this idea, we further performed simulations of FUS-PrLD condensates with different system sizes (by varying the number of protein chains *N*) and calculated their *τ*(*q*) respectively [see Fig. 3(b)]. The resulting curves collapse onto a single master curve with the same exponent of −3.6. For each system, the cutoff depends on the minimum *q*_*min*_ corresponding to the system size (hereafter, we estimated the system size by its linear dimension *L* = 2*π*/*q*_*min*_). Plotting *τ*_*sys*_ against *L* for different system sizes [see Fig. 3(c)], we can see the relationship *τ*_*sys*_ ∼ *L*^3.6^ which has the same form as *τ*_*sys*_ ∼ *ξ*^3.6^. These results demonstrate that the power-law coupling of *τ*(*q*) can be expressed as

**FIG. 3.**
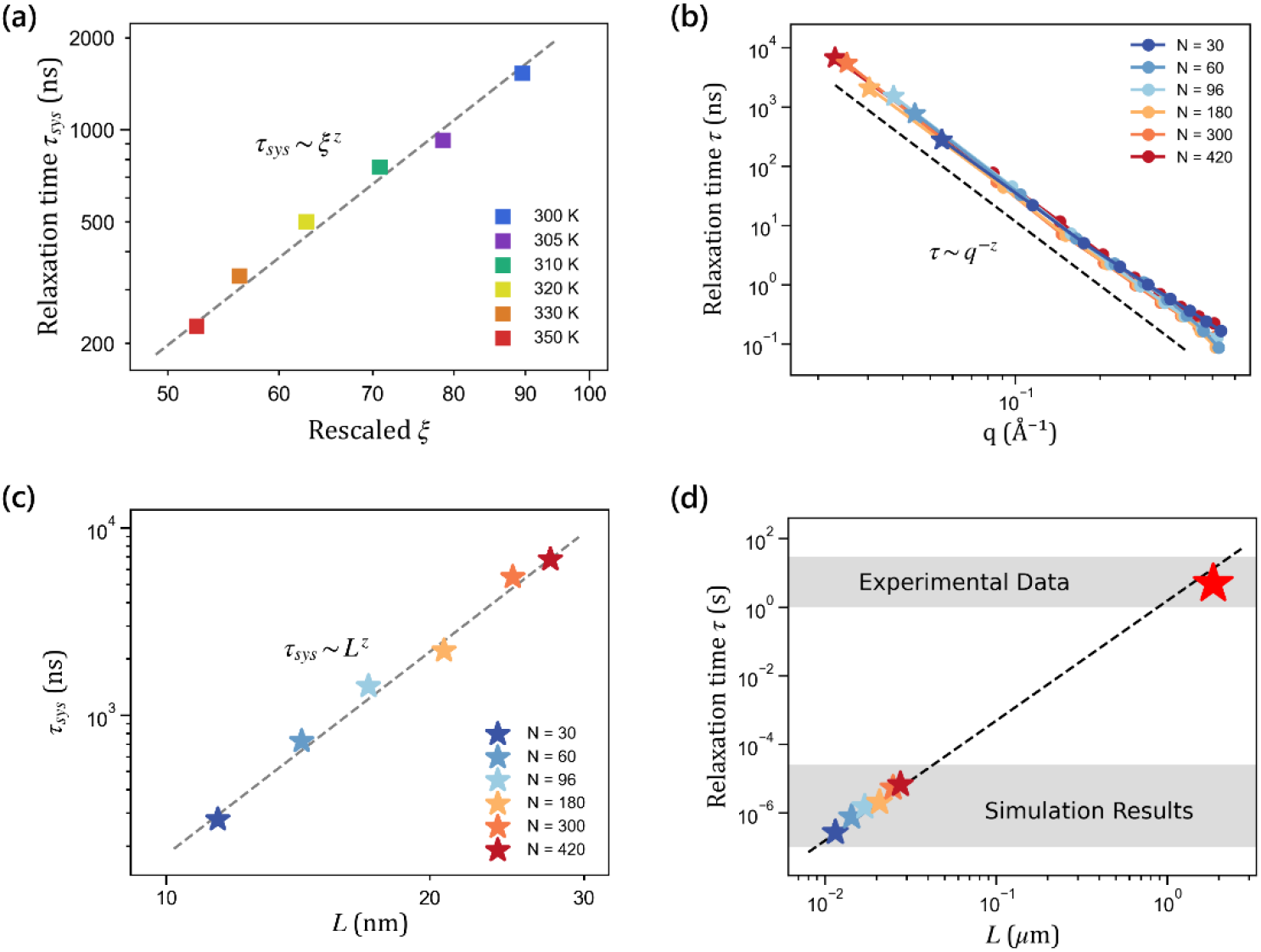
Dynamic scaling of the condensate. (a) Dynamic scaling derived from different temperature systems. The relaxation time of the condensate and the correlation length follow a power-law scaling *τ*_*sys*_ ∼ *ξ*^*z*^, where *z* is the dynamic critical exponent. The dashed grey line represents a fit with *z* = 3.6. (b) As the protein chains *N* in the condensate increases, the system size grows and the relation *τ* ∼ *q*^−3.6^ extends over a wider range. (c) Dynamic scaling obtained from different size systems. *τ*_*sys*_ (star symbols from b) scales with system size as *τ*_*sys*_ ∼ *L*^*z*^, *z* = 3.6. (d) Experimental measurement [36] (red star) aligns with the dynamic scaling prediction from our simulations.

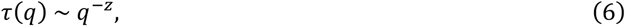

and this relationship observed within a condensate system is essentially the same as the dynamic scaling law *τ*_*sys*_ ∼ *ξ*^*z*^ obtained across systems. Furthermore, the results shown in Fig. 3(b)-3(c) reveal the scale-free nature of the condensate: the spatial range of its dynamic scaling law continues to extend as the system size increases, confirming the point mentioned earlier. Importantly, owing to this scale-free nature, the dynamic scaling law provides a way to bridge findings across scales, e.g., connecting microscopic simulation results with macroscopic experimental data. As shown in Fig. 3(d), an experimental measurement of the FUS-PrLD condensate [36] lies on the *τ*_*sys*_ ∼ *L*^3.6^ scaling, consistent with our simulation prediction.

Notably, the above analysis of the dynamic critical exponent *z* indicates that a consistent value of *z* can be obtained from two independent methods. Now let us examine the relationship between the three critical exponents obtained in the FUS-PrLD condensate. We determined the correlation length critical exponent *v* = 0.26, the dynamic critical exponent *z* = 3.6, and the combined exponent *vz* = 0.93. The values of these exponents satisfy the product relation *vz* = *v* · *z*, which reflects the existence of the following scaling behavior in the condensate:

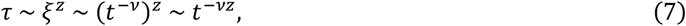

in agreement with critical scaling theory [35]. Taken together, our results demonstrate that the condensate is a critical system and can be characterized by a set of critical exponents.

Given that the dynamic scaling law *τ* ∼ *ξ*^*z*^ provides a comprehensive description of both the static and dynamic scaling behavior of critical systems, the exponent *z* is often used to define their universality classes [5,37]. To investigate whether diverse biomolecular condensates can be characterized by the universality class, we examined several additional condensates: LAF-1 RGG [38], FUS LC [38], ProTα-H1 [39], and ProTα-protamine [40] condensates (all data from published studies). All of these condensates consist of IDPs. The dynamic critical exponent *z* was determined from *τ*(*q*) calculated within each condensate. As shown in Fig. 4(a), the *τ*(*q*) curves for all condensates exhibit power-law scaling, and share similar exponents *z* ≈ 3.6. This result suggests that these IDP condensates likely belong to the same universality class.

**FIG. 4.**
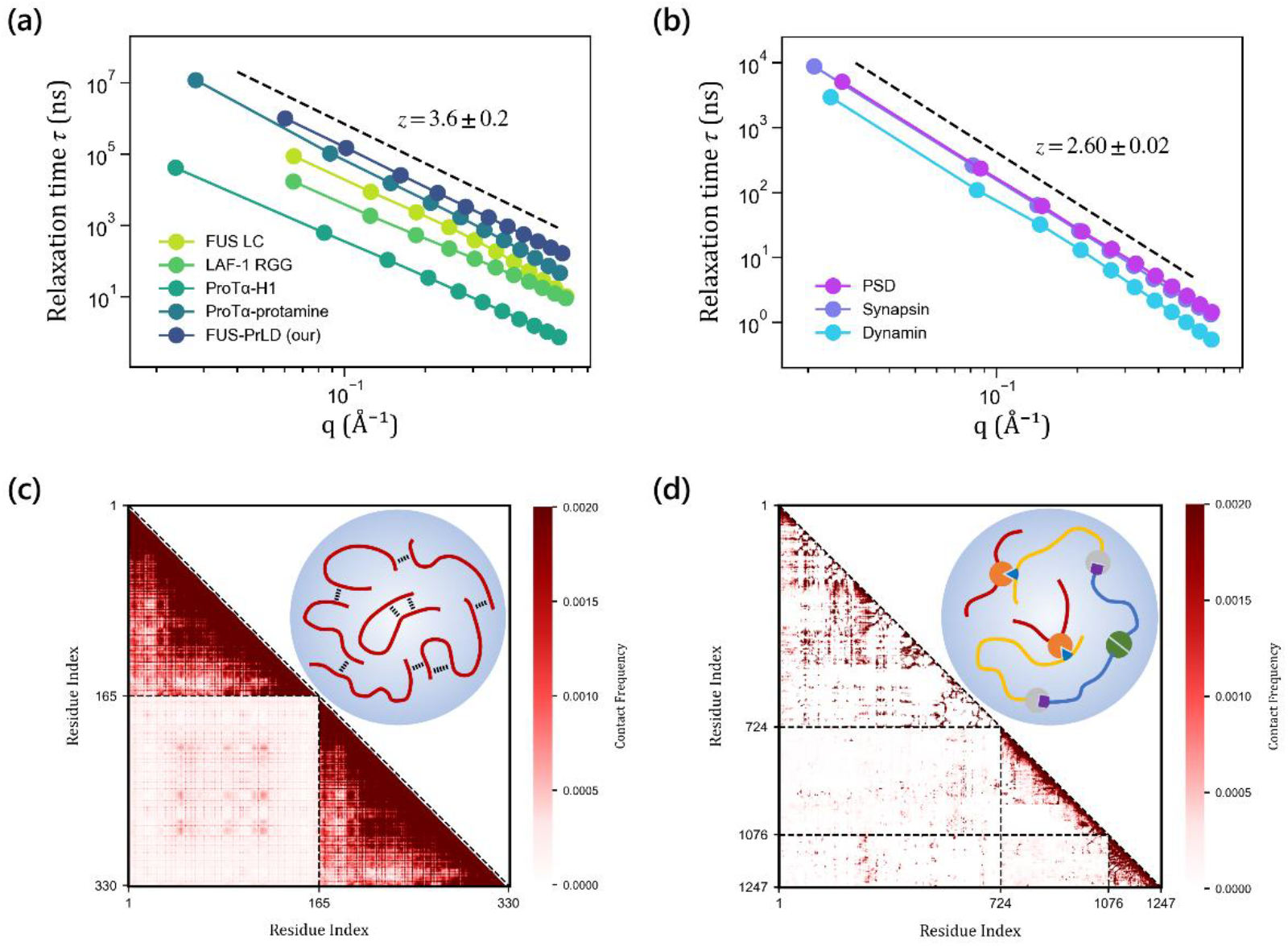
Universality classes of biomolecular condensates. (a) Dynamic scaling of various IDP condensates. They share similar dynamic critical exponents (*z* = 3.6 ± 0.2), suggesting that these systems belong to the same universality class. (b) Dynamic scaling of various MDP condensates. A distinct exponent (*z* = 2.6 ± 0.02) identifies another universality class. (c) Contact map of IDP (FUS-PrLD) condensate. Diagonal and off-diagonal regions correspond to intra- and inter-protein interactions, respectively. (Inset) Schematic of the non-specific interactions within IDP condensate. (d) Contact map of MDP (PSD) condensate. (Inset) Schematic of the specific interactions within MDP condensate.

In addition to IDPs, recent studies have shown that multi-domain proteins (MDPs) can also drive the formation of biomolecular condensates [41,42]. Unlike IDPs whose structures are highly disordered, MDPs contain stable structural domains. We further performed simulations of several MDP condensates: postsynaptic density (PSD), synapsin, and dynamin condensates (see details in the Supplemental Material). As illustrated in Fig. 4(b), the *τ*(*q*) curves of these MDP condensates also follow the power-law relationship *τ*(*q*) ∼ *q*^−*z*^ with similar exponents. However, the dynamic critical exponent here is *z* ≈ 2.6, contrasting with *z* ≈ 3.6 observed in IDP condensates. Thus, MDP condensates can be assigned to a distinct universality class, separate from that of IDP condensates.

The distinction in universality classes between IDP and MDP condensates probably arises from differences in the interaction types within these systems, which depend on the molecular features of their components (i.e., whether containing structural domains). We analyzed interactions in both classes of condensates by computing residue-resolution contact maps. Fig. 4(c) shows the contact map for several proteins in FUS-PrLD condensate. The contacts are sparse and dispersed, reflecting weak non-specific interactions among IDPs. In contrast, the contact map of PSD condensate [see Fig. 4(d)] displays strip-shaped patterns, corresponding to the strong specific interactions between structural domains. Schematic diagrams summarizing the different interaction types in the two classes of condensates are illustrated in Fig. 4(c)-4(d) inset. Notably, in classical physical systems, interaction type is one of the well-established determinants of the universality class of criticality [43,44].

In conclusion, we demonstrated that biomolecular condensates are critical systems, exhibiting various critical phenomena, including scale-free spatiotemporal correlations, divergence of correlation length, critical slowing down, and dynamic scaling law. From these scaling relationships, we derived a set of critical exponents that characterize the behavior of condensates near criticality. Furthermore, we found that condensates composed of different types of proteins belong to two distinct universality classes, which arise from differences in interaction types within condensates.

Our findings open new theoretical and experimental avenues. While we have established a framework for studying the criticality of biomolecular condensates, several fundamental questions remain: First, what is the underlying mechanism of the criticality in condensates? Given the heterogeneity observed within condensates, we propose the “Griffiths phase” as a potential explanation [45] — a concept also used to explain other heterogeneous critical systems such as the brain [46]. Second, what physical principles govern the universality classes of condensates? Renormalization group theory could be introduced in further studies. Moreover, the scale-free properties and scaling behaviors of condensates emerging from criticality, offer a promising path toward bridging simulation and experimental results, thereby enabling meaningful cross‐scale comparisons in the future.

## ACKNOWLEDGEMENTS

We thank Haijun Zhou, Kim Christensen, Hong Qian, Ruoyao Zhang, Mingjie Zhang, Qi Ouyang, Hai Lei, Xiandeng Wu and Xiangze Zeng for constructive comments and helpful discussions. This work was supported by the National Natural Science Foundation of China (NSFC) (Grant Nos. 12175195 and 32371299) and the National Key Research and Development Program of China (Grant No. 2025YFC2311702). We also thank Benjamin Schuler, Robert B. Best and Wenwei Zheng for sharing atomistic simulation trajectories.

## AUTHOR CONTRIBUTIONS

H.S. and J.L. conceptualized and designed the project. H.S., G.H. and X.Z. performed the simulations. H.S. and X.W. conducted the data analysis. H.S. wrote the manuscript. All authors discussed and revised the manuscript.

## DATA AVAILABILITY

All data needed to evaluate the conclusions in the paper are present in the paper and/or the Supplementary Materials. Additional data related to this paper are available from the corresponding authors upon request.

