## Supplementary Materials for "Critical Scaling Laws and Universality Classes in Biomolecular Condensates"

TABLE OF CONTENTS

I. Simulation Details
Section A: All-atom Molecular Dynamics Simulations
Section B: Coarse-grained Molecular Dynamics Simulations

II. Computational Methods
Section A: Power Spectrum
Section B: Structure Factor
Section C: Intermediate Scattering Factor
Section D: Linear Dimension of Condensates
Section E: Correlation Length
Section F: Contact Map
Section G: Data from Experiment

III. Supplementary Data

Additional References

I. Simulation Details

A. All-atom Molecular Dynamics Simulations

The prion-like domain (PrLD) of Fused in Sarcoma (FUS) is a typical intrinsically disordered protein (IDP). It can undergo liquid-liquid phase separation (LLPS) and form biomolecular condensates. We investigated the condensate composed of FUS-PrLD using all-atom molecular dynamics (MD) simulations. The FUS-PrLD condensate contained 48 protein chains. We constructed a slab geometry with box dimensions of 14 × 14 × 35 $nm^3$ (X × Y × Z) as the initial configuration for the condensate simulation (Fig. S1), following the steps in previous studies [1].

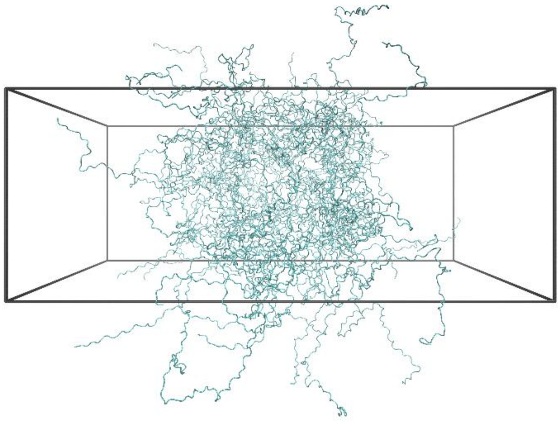

FIG. S1. The initial configuration of the FUS-PrLD condensate with a slab geometry for the all-atom simulation.

MD simulations were performed using GROMACS software package (version 2023.5 or later). Amber a99SB-disp force field with the modified TIP4P-D water model (a99SB-disp water model) was used [2]. The slab box with initial condensate configuration was solvated by TIP4P-D water model, and sodium and chloride ions were added to neutralize the system and mimic physiological conditions (150 mM NaCl). The system was energy-minimized and then equilibrated first under NVT ensemble at 300 K for 50 ns, followed by NPT ensemble at 1 atm for 1100 ns. Temperature and pressure were maintained using velocity-rescaled thermostat and Parrinello-Rahman barostat, respectively. After the extensive equilibration, a 3000-ns production simulation was conducted in the NVT ensemble at 300 K.

The all-atom simulations employed a leapfrog integrator with a 2-fs time step. The periodic boundary conditions (PBC) were applied in all directions. A 1.2-nm cutoff was used for both short-range electrostatic interactions and van der Waals (VdW) interactions. The particle mesh Ewald (PME) method was used for long-range electrostatic interactions. The Verlet cut-off scheme was employed for neighbor searching. The covalent bonds involving hydrogen atoms were constrained by the LINCS algorithm. The system coordinates were saved every 50 ps.

The all-atom MD simulation trajectories of additional condensates were obtained from the available data shared by the authors of the published studies. These condensates are: FUS LC and LAF-1 RGG condensates (data from Ref. [1]), ProTα-H1 condensate (from Ref. [3]), ProTα-protamine condensate (from Ref. [4]).

B. Coarse-grained Molecular Dynamics Simulations

1. General CG simulation settings

Various condensates were studied using coarse-grained (CG) molecular dynamics simulations, taking advantage of the high computational efficiency. These CG simulations employed the following general settings. GROMACS software package (version 2023.5 or later) was used. We employed the MARTINI 3.0 force field [5]. The simulation boxes were solvated by MARTINI water with the salt concentration of 150 mM NaCl. We used TQ5 bead types for $Na^+$ and $Cl^-$ ions. The systems were energy-minimized and then equilibrated, first under an NVT ensemble for 2 ns, followed by an NPT ensemble at 1 atm for 500 ns. Temperature was maintained using velocity-rescaled thermostat with coupling constant 1 ps. Pressure was maintained using Parrinello-Rahman barostat with coupling constant 4 ps. The leapfrog algorithm was used to integrate the equations of motion with a time step of 20 fs. The PBC were applied in all directions. The Verlet neighbor search algorithm was used to update the neighbor list. A cutoff distance of 1.1 nm was used for the van der Waals interactions. Coulomb interactions were treated by a reaction-field with dielectric constant 15 and 1.1 nm cutoff.

2. FUS-PrLD condensates at different temperatures

CG simulation systems for FUS-PrLD condensates at different temperatures (300 K, 305 K, 310 K, 320 K, 330 K, and 350 K) were constructed as follows. All-atom coordinates of FUS-PrLD protein chains with diverse conformations were converted to coarse-grained coordinates using Martinize2. A total of 96 FUS-PrLD chains were inserted randomly inside a cubic box of 21 nm edge, then expanded to a larger box of 32 nm, and subsequently solvated. A 1000-ns simulation was performed on this system to form a droplet (spherical condensate; Fig. S2). The resulting droplet was then equilibrated at each target temperature for 500 ns. The final equilibrated configurations were used as the starting configurations for 5000-ns production simulations at their respective temperatures. To reproduce a reasonable droplet of FUS-PrLD condensate, the protein-water interaction was enhanced by 0.1 kJ/mol (~2%) relative to the unmodified MARTINI 3.0 force field, as suggested in previous work [6]. This adjustment was implemented by adding water-bias between protein and water beads using Martinize2.

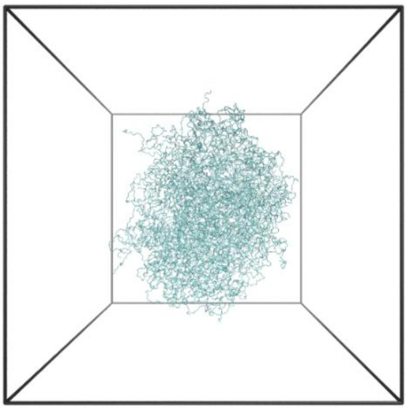

FIG. S2. The configuration of FUS-PrLD condensate (droplet) for the coarse-grained simulations.

We examined the critical behaviors of FUS-PrLD condensates obtained from all-atom and CG simulations under the same conditions (300 K, 1 bar, 150 mM NaCl). The scale-free spatiotemporal correlations of the all-atom system were shown in Fig. 1 in the main text. Similarly, scale-free correlations were observed in the CG system (Fig. S3). The dynamic scaling laws for both systems were shown in Fig. S4. Both systems exhibit a similar dynamic critical exponent ($z \approx 3.6$). The difference in their relaxation time is primarily caused by the different resolutions of the two simulation methods: the Martini CG simulation represents residues with a reduced number of beads (typically 1-6 beads per residue), whereas the all-atom simulation includes full atomic details. This difference in resolution influences the frictions among residues and their dynamics within condensates, thereby leading to the difference in relaxation time. Nevertheless, despite the different simulation methods, the same scale-free correlations, scaling behaviors and dynamic critical exponent were observed in both systems. This can also reflect that the universal features of critical systems are largely independent of their specific microscopic details.

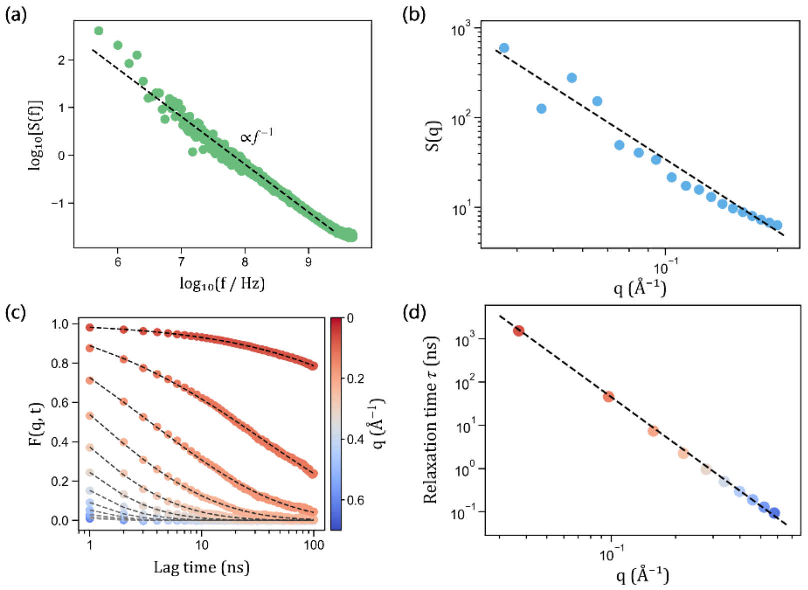


FIG. S3. Scale-free spatiotemporal correlations in CG simulation of the FUS-PrLD condensate. (a) $1/f$ power spectrum. (b) Power-law structure factor. (c) ISF curves for different $q$. (d) Dynamic scaling within condensate.

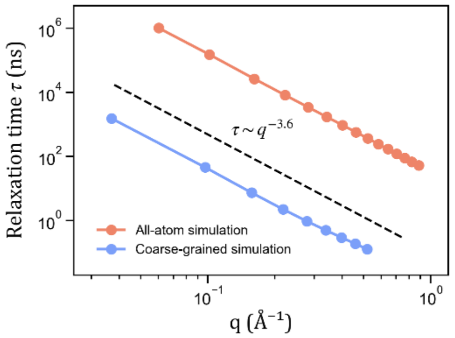


FIG. S4. The dynamic scaling within the FUS-PrLD condensate for both all-atom and CG simulations. The exponents obtained from the two simulation methods are consistent: $z \approx 3.6$.

### 3. FUS-PrLD condensates with different sizes

CG simulation systems for FUS-PrLD condensates of varying sizes were constructed, containing protein chains $N = 30, 60, 96, 180, 300, 420$. To construct spherical condensates with different sizes, $N$ chains of FUS-PrLD were inserted randomly inside small cubic boxes. These boxes were then expanded to larger boxes and solvated. The corresponding box dimensions were provided in Table S1. Each system was simulated for 1000 ns to form spherical condensates. The condensates were then equilibrated under the NPT ensemble for 500 ns. These final equilibrated configurations were used as the starting configurations for 2000-ns production simulations under 300 K, 1 bar, 150 mM NaCl. The configurations of condensates with different sizes were represented in Fig. S5.

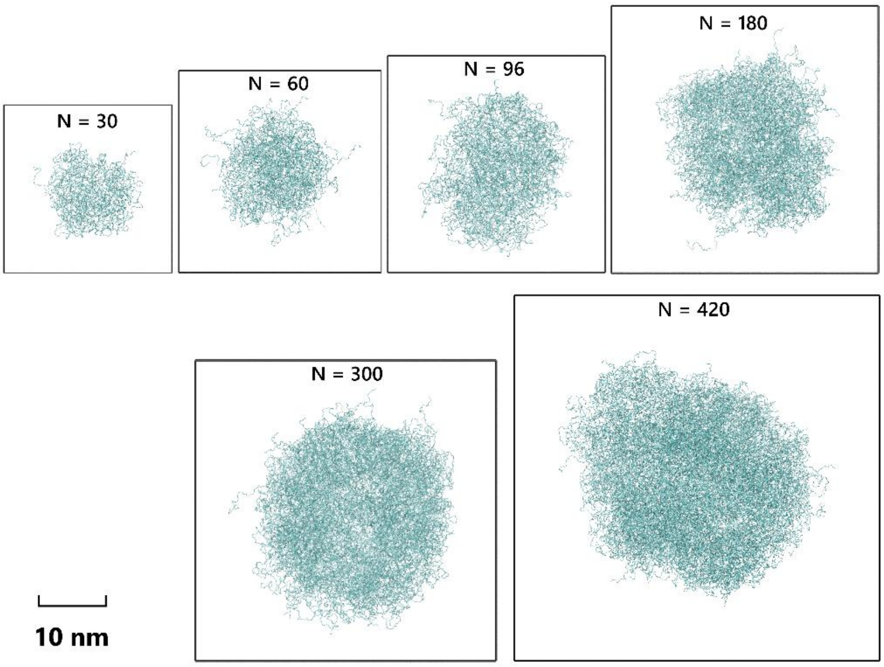


FIG. S5. Configurations of FUS-PrLD condensates of different sizes (front view) from CG simulations.

| Protein Chains $N$ | Cubic Box Length (nm) |
|---|---|
| 30 | 25 |
| 60 | 30 |
| 96 | 32 |
| 180 | 40 |
| 300 | 45 |
| 420 | 55 |

TABLE S1. Simulation box parameters for the CG simulations of the FUS-PrLD condensates with different sizes.

4. MDP condensates

We also performed CG simulations of three multi-domain protein (MDP) condensates. MDPs contain structural domains, linked by intrinsically disordered regions (IDRs). Three MDP condensates studied were postsynaptic density (PSD), synapsin condensate, and dynamin condensate.

The PSD of neuronal synapses is a specialized cellular condensate essential for synaptic transmission, memory formation, and storage [7–9]. Our PSD system comprised three scaffold proteins: PSD-95 (UniProt: P78352-1; aa 1-724), GKAP (UniProt: Q9D415-1; aa 328-421+916-992), SHANK3 (UniProt: Q4ACU6-1; aa 533-665+1294-1426+1645-1730), as previously reported. The PSD condensate contained 20 chains of each protein.

Synapsin can form condensate via LLPS and intersectin (a protein with multiple SH3 domains) enhances the stability and kinetics of condensate formation [10–12]. This condensate can capture synaptic vesicles (SVs) at the presynaptic terminal, as a reservoir that ensures neurotransmitter release. Our synapsin condensate included synapsin (UniProt: O88935-1; aa 1-706) and intersectin (UniProt: Q15811-1; aa 740-1214), with 50 chains of each protein.

Fast endophilin-mediated endocytosis (FEME) is a clathrin-independent endocytosis (CIE) pathway. LLPS plays a crucial role in the formation of endophilin-rich condensates that serve as initiation sites for FEME, and these condensates further recruit dynamin to promote membrane scission [13,14]. Our dynamin condensate consisted of dynamin (UniProt: Q05193-1; aa 1-864) and endophilin (UniProt: Q62420; aa 1-352), with 25 chains of each protein.

The structures of these MDPs were predicted by AlphaFold3 [15]. The predicted structures were then energy-minimized, equilibrated, and relaxed using all-atom simulations. After relaxation, all-atom coordinates of MDPs were converted to coarse-grained coordinates using Martinize2. An intradomain elastic network model with harmonic potentials of 700 $kJ\cdot mol^{-1}\cdot nm^{-2}$ was applied to keep the structural domain intact. The -scFix option in Martinize2 was employed to restrict side chain orientation in the structural domain. The water-bias was added to IDR regions using Martinize2. As demonstrated previously [16], Martini force field underestimates inter-domain interactions in aqueous solution, leading to the rapid and irreversible dissociation of oligomers in the beginning of the simulations. To address this, we adopted an approach analogous to the Gō model as suggested: virtual sites with an enhanced intermolecular potential of ~10 kJ·mol$^{-1}$ were applied to the inter-domain interactions of the MDPs studied here, using a Python script provided in the previous work [16]. A 1500-ns CG simulation was performed for each MDP system to form condensate. The production simulations were then performed under 300 K, 1 bar, 150 mM NaCl for 2000 ns.

The all-atom structure of PSD-95 and the CG simulation snapshot of PSD condensate were shown in Fig. S6. Similar to the FUS-PrLD condensate, the PSD condensate also exhibits scale-free correlations (Fig. S7).

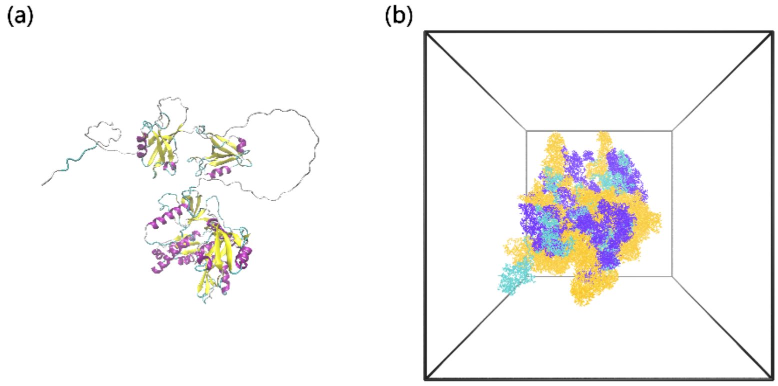


FIG. S6. Configuration of PSD condensate. (a) The all-atom structure of PSD-95 with structural domains and IDRs. (b) The configuration of PSD condensate containing PSD-95 (orange), GKAP (cyan) and SHANK3 (purple).

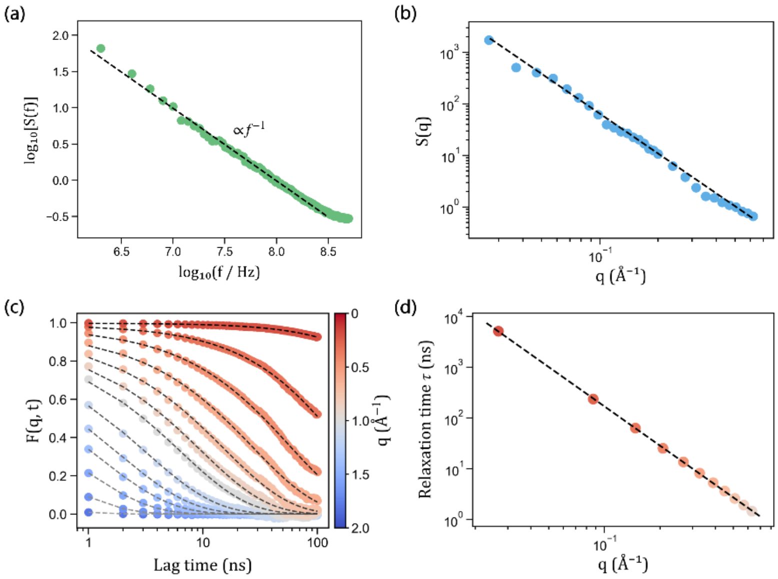


FIG. S7. Scale-free spatiotemporal correlations in CG simulation of the PSD condensate. (a) $1/f$ power spectrum. (b) Power-law structure factor. (c) ISF curves for different $q$. (d) Dynamic scaling within condensate.

### II. Computational Methods

#### A. Power Spectrum

The temporal correlation can be characterized by the shape of the power spectrum. We calculated the power spectrum of local residue density fluctuations. A spherical region with a radius of 4 nm was defined around the center of mass of the condensate, and the number of residues within this region was computed over time. The local residue density fluctuations, $n(t)$, was defined as the time-dependent number of residues within the region divided by its volume. Its power spectrum $S(f)$ was then calculated by Fourier transforming the autocorrelation function of $n(t)$,

$$S(f) = \int \langle n(t_0 + t) n(t_0) \rangle e^{-2\pi i f t} dt, \tag{S1}$$

where the angular brackets $\langle \cdot \rangle$ represent an average over all times $t_0$.

#### B. Structure Factor

The structure factor $S(q)$ can reflect the spatial correlation of the condensate in the reciprocal space. Particle positions were extracted from the simulation trajectories. The vector $\vec{r}_j(t)$ represents the position of $j$-th atom (for all-atom simulations) or $j$-th bead (for CG simulations) at time $t$. The structure factor of the condensate was then calculated as

$$S(q) = \frac{1}{N} \langle \left| \sum_{j=1}^{N} e^{-i\vec{q}\cdot\vec{r}_j(t)} \right|^2 \rangle. \tag{S2}$$

Here, the angular brackets $\langle \cdot \rangle$ represent an average over all times $t$, $\vec{q}$ is the wavevector with the corresponding wavenumber $q$. For each condensate conformation (at time $t$) and for each wavenumber $q$, $S(q)$ was evaluated by averaging over 5 randomly oriented $\vec{q}$.

#### C. Intermediate Scattering Factor

To examine the coupling between spatial and temporal correlations, we also considered the intermediate scattering function (ISF) $F(q,t)$, which is a time-dependent correlation function. Notably, the static $S(q)$ corresponds to the ISF at time zero, i.e., $S(q) = F(q,0)$. The decay of $F(q,t)$ with time provides direct insight into the relaxation dynamics of the system. The ISF of the condensate was calculated as follows:

$$F(q,t) = \frac{1}{N} \langle \sum_{i=1}^{N} \sum_{j=1}^{N} e^{-i\vec{q}\cdot\vec{r}_i(t_0+t)} e^{i\vec{q}\cdot\vec{r}_j(t_0)} \rangle, \tag{S3}$$

where the angular brackets $\langle \cdot \rangle$ represent an average over all times $t_0$, $t$ is the lag time, and $\vec{q}$ is the wavevector with the corresponding wavenumber $q$. For each wavenumber $q$, the corresponding relaxation time $\tau(q)$ was obtained by fitting the ISF decay to a stretched exponential function

$$F(q,t) = Ae^{-\left[\frac{t}{\tau(q)}\right]^{\beta(q)}}, \tag{S4}$$

where $A$ is the amplitude, $\tau(q)$ is the relaxation time, and $\beta(q)$ is the stretching exponent. For convenience, the ISF of condensates was calculated with $\vec{q}$ aligned along the simulation box z-axis. As shown in Fig. S8 for FUS-PrLD condensate, the $\tau(q)$ curves calculated using $\vec{q}$ along the x-, y-, z-axis all exhibit power-law relation and the similar exponents. This indicates that the choice of wavevector direction has minimal effect on the measured $\tau(q)$.

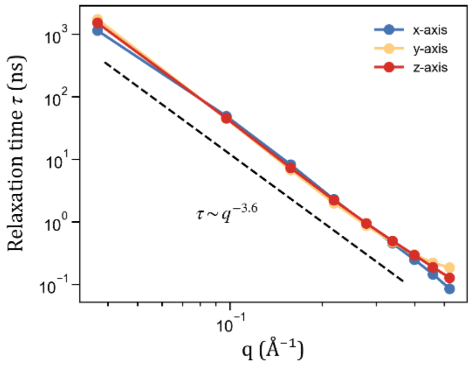


FIG. S8. $\tau(q)$ curves obtained using wavevector $\vec{q}$ along the x-, y-, z-axis of the simulation box, respectively. These curves exhibit the same behavior.

D. Linear Dimension of Condensates

With the scale-free properties of the condensates, the spatial range exhibiting dynamic scaling expands as the system size grows. To describe the system size of the condensates, the linear dimension $L$ was used. For spherical condensate droplets, $L$ was estimated by the calculation of the radius of gyration of the condensate as

$$L \approx 2R_g. \tag{S5}$$

$R_g$ was calculated using the formula:

$$R_g = \sqrt{\frac{\sum_i m_i |\vec{r}_i|^2}{\sum_i m_i}}, \tag{S6}$$

where $m_i$ is the mass of the $i$-th atom or bead, and $\vec{r}_i$ is its position with respect to the condensate's center of mass. For slab-geometry condensates (where biomolecules occupy the xy-plane and $\vec{q}$ for ISF calculation is along the z-axis, perpendicular to the xy-plane), $L$ was taken as the box length along x-axis (the same box length along x- and y-axis in our simulations). The linear dimension $L$ in real space corresponds to a minimum wavenumber in reciprocal space,

$$q_{min} = \frac{2\pi}{L}. \tag{S7}$$

The system relaxation time of the whole condensate, $\tau_{sys}$, was then defined as relaxation time $\tau(q)$ at $q = q_{min}$:

$$\tau_{sys} = \tau(q_{min}). \tag{S8}$$

E. Correlation Length

We computed the velocity-velocity correlation function analogous to the algorithm used in studies of flocking birds [17]. This correlation function reflects whether the residues within condensates move coherently. To estimate the velocity of the residue, we first extracted the center-of-mass coordinates of all residues at neighboring frames $t$ and $t+1$ from the trajectory. The residue velocity was calculated as

$$\vec{v}_i = \frac{\vec{x}_i(t+1) - \vec{x}_i(t)}{\Delta t}, \tag{S9}$$

where $\vec{x}_i(t)$ is the coordinate of the $i$-th residue at frame $t$, and $\Delta t$ is the time interval between neighboring saved trajectory frames. To remove the global translational motion of the condensate, the center-of-mass velocity of the condensate $\bar{v}$ was subtracted from the residue velocity

$$\vec{u}_i = \vec{v}_i - \bar{v}, \tag{S10}$$

to obtain the relative velocity of residue. Then the correlation for a pair of residues $i$ and $j$ is then defined as the dot product of their relative velocities:

$$C_{ij} = \vec{u}_i \cdot \vec{u}_j. \tag{S11}$$

With the consideration of correlations of all residue pairs throughout the condensate, the correlation function was defined as follows,

$$C(r) = \frac{\sum_{i<j}^{N} \frac{C_{ij}}{\sqrt{C_{ii}C_{jj}}} \delta(r - r_{ij})}{\sum_{i<j}^{N} \delta(r - r_{ij})}, \tag{S12}$$

where $\delta(r - r_{ij})$ is the Dirac delta function, and the mutual distance $r_{ij}$ of residue pairs was computed as their center-of-mass distance. Fig. S9(a) shows the correlation function $C(r)$ for the FUS-PrLD condensate. Its profile is similar to those reported in previous studies. As established previously [17], the correlation function takes the form

$$C(r) = \frac{1}{r^{\gamma}} f\left(\frac{r}{L}\right), \tag{S13}$$

where $f(r/L)$ is a dimensionless scaling function cut off by system size $L$. This expression indicates that $C(r)$ exhibits a power-law dependence on $r$, but is strongly influenced by finite-size effect. As shown in Fig. S9(b), $C(r)$ of the condensate follows the power law, with deviations at large $r$ (the tail of the curve) reflecting finite-size effects. Given that the difference in the size of condensates at different temperatures is not significant, we estimated the correlation length of them as follows to circumvent finite-size limitation: the power-law region of $C(r)$ was fitted to $C(r) \sim r^{-\gamma}$ [Fig. S9(c)], and the rescaled correlation length $\xi$ was then defined as the value of $r$ at which the extrapolated power-law fit decays to 0.01 [Fig. S9(d)].

To verify the reasonableness of the critical exponent $\nu$ obtained from this rescaled $\xi$ in the relationship

$$\xi \sim t^{-\nu}, \tag{S14}$$

we employed an independent consistency check. This check used the dynamic critical exponent $z$, which was independently determined via two methods (Fig. 3). Using the scaling relation $\tau_{sys} \sim \xi^z$ with $z = 3.6$, we plotted the $L^{-1}\tau_{sys}^{1/z}$ versus $t$, where $L$ is the linear dimension of the condensate and $t$ is the reduced temperature. As shown in Fig. S10, the power-law relation

$$\tau_{sys}^{1/z} \sim Lt^{-\nu} \tag{S15}$$

can be observed. Notably, the value of $\nu$ obtained from this scaling plot is 0.26, identical to the value derived from the rescaled $\xi$ in Fig. 2(b). Thus, the value of $\nu = 0.26$ is validated by this independent scaling analysis, confirming the reasonableness of both the rescaled $\xi$ and the derived critical exponent $\nu$.

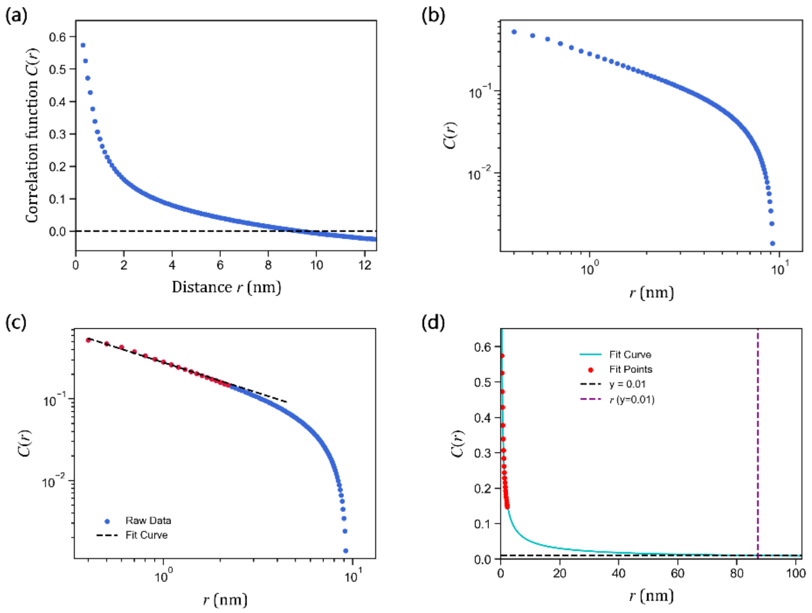


FIG. S9. The correlation function $C(r)$. (a) Correlation function of the FUS-PrLD condensate at 300 K. As $r$ increases, the correlation decays from its maximum and crosses zero. (b) Log-log plot of the correlation function. (c) Fitting of the power-law region of the correlation function. (d) The extrapolation of fit curve. The rescaled $\xi$ was estimated as $r$ at which correlation function decays to 0.01.

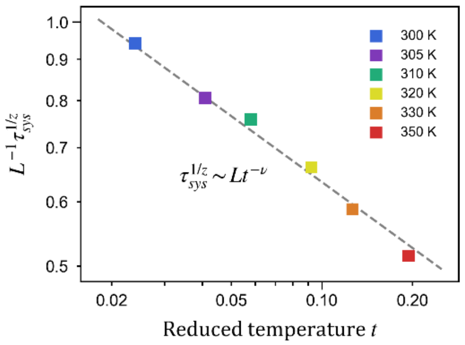


FIG. S10. Additional estimation of the critical exponent $\nu$ for the correlation length. The fitted value $\nu = 0.26$ matches that obtained in Fig. 2(b).

### F. Contact Map

The contact maps for both FUS-PrLD (IDP) condensate and PSD (MDP) condensate were calculated using the trajectories from CG simulations. A contact between two residues was defined if the minimum distance between any of their corresponding beads was less than 0.6 nm. The contacts of neighboring residues along the sequence (i.e., the contacts between residue $i$ and $i \pm 1, \pm 2$) were excluded. After the calculation of residue contacts, the contact map can be obtained. The square blocks along the diagonal represent intra-molecular contacts, averaged over all contacts within the same protein type. The rectangular blocks off the diagonal represent inter-molecular contacts, averaged over all pairs of proteins that were in contact.

### G. Data from Experiment

Dynamic scaling law can bridge the gap between microscopic simulation results and macroscopic experimental data. We extracted an experimental data point from the fluorescence recovery after photobleaching (FRAP) experiment of FUS LC droplet condensates (reported in Fig. 2 of Ref. [18]). We estimated the relaxation time $\tau$ and the spatial length $L$ of the droplet as follows: The relaxation time was estimated by the fluorescence half-time ($T_{1/2}$) of the droplet presented in Fig. 2(d), i.e., $\tau \approx T_{1/2} \approx 5\ s$. The spatial length $L$ was estimated by the radius of the bleached region presented in Fig. 2(a), i.e., $L \approx 1.85\ \mu m$.

### III. Supplementary Data

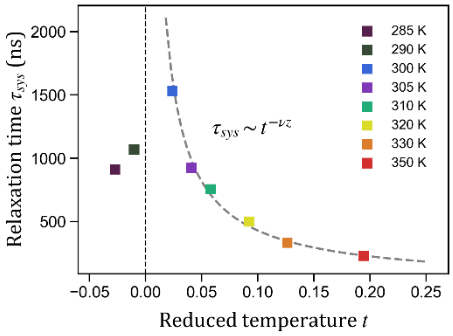


FIG. S11. Divergence of the condensate behavior near the critical point. The critical temperature was estimated as the room temperature, i.e., $T_c \approx 293\ K$. The relaxation time of the condensate exhibits a power-law dependence on reduced temperature $t = T/T_c - 1$: $\tau_{sys} \sim t^{-\nu z}, t \to 0^+$. Additionally, we performed two CG simulations (290 K, 285 K) below $T_c$ and observed a corresponding decrease in $\tau_{sys}$.
